# CONTROL OF *OROBANCHE CUMANA* WALLR: RESEARCH ON SUSTAINABILITY OF SUNFLOWER HYBRIDS AND STRATEGIES FOR PARASITE PROTECTION

**DOI:** 10.1101/2024.10.12.617968

**Authors:** Serhii Khablak, Lesia Bondareva, Mykola Dolia, Yaroslav Blume, Tetiana Tymoshchuk, Ivan Mrynskyi, Natalia Hrytsiuk, Valentyn Spychak

## Abstract

The work is devoted to solving an important scientific task of determining the spread and harmfulness of broomrape, its racial composition, and developing measures to protect sunflower from this parasite. The aim of the research was to determine the resistance of sunflower hybrids to the broomrape parasite. During the research, a vegetation experiment was used to determine the racial composition of the parasite and the resistance of different sunflower hybrids to it. Sunflower hybrids were evaluated for resistance to broomrape in soil culture using a modified method and the roll method of seed germination. From the northern Steppe of Ukraine, broomrape is actively moving to the central, northern and western regions of the country. The practical value of the work is to determine the resistance of sunflower hybrids to broomrape. Differentiation of sunflower hybrids grown by resistance to the parasite was carried out. It has been observed that the broomrape population exhibits a significant level of virulence capable of overcoming the immunity found in the most resilient foreign-bred hybrids resistant to E, F, and G races of this parasite. The appearance of highly aggressive new races of broomrape (E, F, G, and H) in the Forest-Steppe and Polissya regions underscores the crucial necessity to address the challenge of developing breeding material resistant to these novel races of the parasitic plant. This entails an in-depth exploration of the cellular and molecular mechanisms underlying sunflower resistance to the pathogen. The widespread buildup of parasite races E, F, G, and H in sunflower crops is linked to the disruption of crop rotations and the extensive cultivation of hybrids of this crop. These hybrids are primarily resistant to races 5 (E) and 6 (F) of the parasite.

The results of the research can be used in the region’s farms for successful protection against broomrape, as well as in breeding programs to create sunflower hybrids resistant to new races of the parasite and corn hybrids that cause better germination of seeds of this pathogen in the soil and their death by their root secretions.

## INTRODUCTION

In Ukraine, at the beginning of the XXI century, a significant increase in the areas under sunflowers contributed to the spread of the parasite *Orobanche cumana* Wallr. *Orobanche cumana* is a parasitic plant that infects the plant’s underground system and absorbs water and nutrients from it. The solution of an important scientific task to determine the features of the development of sunflower broomrape and the development of measures to limit its harmfulness in the conditions of forest-steppe and Polissya requires theoretical justification and new practical approaches, which has ensured the priority and relevance of this research.

A parasitic plant is a flowering plant that establishes a morphological and physiological connection with a host (another plant) through a modified root known as haustoria. Among the 270 genera of parasitic plants, only around 25 are deemed detrimental to agriculture and forestry, classifying them as weeds. Notably, the most destructive root parasitic weeds belong to the genera *Orobanche* and *Phelipanche*, commonly referred to as broomrape, as well as *Striga*. All these belong to the family *Orobanchaceae* (Vurro et al., 2019; Shatkovskyi et al., 2022).

According to Albanova et al. (2023) parasitic flowering plants, such as members of the family *Orobanchaceae* and *Cuscuta,* pose a serious threat to agricultural crops. Their methods of parasitism are to extract water and nutrients from their hosts, which can significantly affect the vegetative growth and yield of plants. Key challenges in controlling these parasites include seed longevity and in-soil stability, as well as difficulties in using selective herbicides. Thus, it is important to develop control methods that take these features into account. The selection of resistant varieties is a promising approach. This approach may include studying the molecular mechanisms of host resistance and inheriting these characteristics. Although this approach is not widely used due to limited knowledge of molecular mechanisms, its development may be an important area for reducing crop losses due to parasitism of these plants. It is important to continue research in this direction and develop effective strategies for managing parasitic flowering plants in agriculture.

The development of sunflower varieties resistant to parasitic plants like *O. cumana* necessitates the discovery and identification of novel resistance genes. According to Calderón-González et al. (2023), an analysis of the genetic structure involving 104 sunflower germplasm samples revealed the presence of two primary groups. Utilizing optimized methods based on the general linear model (GLM) and mixed linear model (MLM), researchers identified 14 SNP markers significantly associated with resistance to broomrape. The majority of these marker-trait associations were located on chromosome 3, clustered within two distinct genomic regions. Additional associations were discovered on chromosomes 5, 10, 13, and 16. Calderón-González et al. (2023) thoroughly investigated and discussed candidate genes within major genomic regions linked to broomrape resistance. Specifically, two noteworthy SNPs on chromosome 3, associated with the EFR and FGV races, were identified in two closely related SWEET sugar transporter genes. The study’s findings affirm the role of certain QTL genes in sunflower resistance to broomrape and unveil new candidates that could play a crucial role in developing enduring resistance to this parasitic weed in sunflower.

The study conducted by Aly et al. (2021) underscores the challenge posed by the absence of new reservoirs of resistance, limiting our capacity to effectively manage emerging and more potent broomrape variants. Given that most crops lack efficient methods to control parasites, there is a pressing need for innovative biotechnological solutions. Recent advancements have revealed the efficacy of various strategies employing regulatory RNA molecules, the CRISPR/Cas9 system, and T-DNA inserts in developing resistance against parasitic weeds. In the past years, significant strides have been taken in decoding the plant genome and understanding its functions, including those of parasitic weeds. Nevertheless, further research is essential to deepen the foundation of biotechnological strategies for instilling host resistance to root parasitic weeds. Gene silencing and editing tools should be harnessed to target critical processes in the host-parasite interaction, such as biosynthesis and strigolactone signaling, haustoria development, as well as the degradation and penetration of the host cell wall.

Currently, the study of cellular and molecular mechanisms of sunflower resistance to broomrape is an urgent area for creating new hybrids with high resistance to this pathogen. Access to the genetic resources of sunflower plasma is important for breeders working on creating resistant sunflower hybrids to broomrape. Recently, a publicly available and interactive database of sunflower phenotypes called HelianTHOME has been created (http://www.helianthome.org). This database covers a wide range of sunflower samples, including both wild and cultivated forms of crop. In addition, the database is saturated with external genomic data and results of association studies at the genome level. HelianHOME is predicted to expand with the advent of new knowledge and resources (Bercovich et al., 2022).

According to Chander et al. (2022) recent advances in large-scale DNA sequencing and high-performance screening techniques have significantly changed the approach to crop selection. In the modern world, reverse genetics approaches have become a key area of research in a large number of crops, including sunflower. The adoption of innovative molecular methodologies, notably EcoTILLING within the broader framework of TILLING, has enabled the utilization of naturally occurring genetic variations. This approach involves the identification and utilization of induced or pre-existing genetic changes to facilitate the development of new plant varieties.

Cuccurullo et al. (2022) collected updated information on the genetic and molecular mechanisms underlying host-parasite interactions. While studying the germ plasmas of some species, they have identified natural sources of resistance, and the use of artificial mutagenesis allowed for additional variability. Recent advancements in the genomics of both parasitic plants and their hosts, coupled with emerging breeding technologies, lay the foundation for future progress. As highlighted by Cuccurullo et al. (2022), the integration of diverse genetic mechanisms of resistance, especially interventions targeting various stages of parasite development, along with the incorporation of innovative agronomic management techniques, holds the potential to establish an effective and enduring strategy for pathogen control.

Regrettably, as of now, there is a lack of reported research on the epigenetic mechanisms associated with sunflower resistance. So far, only a handful of researchers have used the sunflower genome sequence in their molecular studies, especially in terms of interaction with broomrape. The development of methods and powerful statistical tools for analyzing big data can be effectively used to study the mechanisms of interaction between sunflower and broomrape, as well as to reveal the ways of crop resistance.

Unfortunately, so far there have been no reports of studying the epigenetic mechanisms of sunflower resistance. According to Cvejić et al. (2020), considering them as a new area of research, it would be important to investigate the effect of epigenetic mechanisms on sunflower resistance, in particular, taking into account the role of DNA methylation in regulating seed germination of *Phelipanche ramosa* during conditioning and control of PrCYP707A1 expression, which depends on strigolactone.

As highlighted by Aly et al. (2021), the highly effective CRISPR-Cas9 technique has been successfully employed for the mutagenesis of the CCD8 (Carotenoid Cleavage Dioxygenase 8) gene, a pivotal gene in strigolactone biosynthesis. This approach has led to the development of tomato lines exhibiting resistance to *Phelipanche aegyptica*. The introduction of new gene editing techniques may pose challenges for sunflower breeding, in particular due to the difficulties associated with the plant regeneration process and the limited number of transgenic regenerants obtained in one experiment. Thus, the first stage of using modern gene editing techniques involves developing an improved transformation framework that could contribute to the development of long-term resistance to broomrape in sunflower.To analyze the evolutionary processes of contagion and develop effective control strategies, it is important to gain a deep understanding of the distinctive features of different races and their pathogenic effects.

According to Duca et al. (2020), the application of simple sequence repeat (SSR) and inter-simple sequence repeat (ISSR) markers has facilitated the identification of specific amplicons capable of distinguishing between different races. The study by Duca et al. (2020) also demonstrated that genes such as PME, PGU, PRX, and CHS, responsible for encoding host invasion-related enzymes like pectinmethylesterase, polygalacturonase, peroxidase, and chalconsintase, respectively, were activated in race H as compared to E at various developmental stages, affirming their role in the pathogenicity of broomrape. Proteomic analysis unveiled 19 differentially expressed proteins (DEP), comprising 6 specific to race E and 13 specific to race H. Among these DEPs are respiratory enzymes, cell wall-modifying enzymes, stress-related proteins, and other proteins known for their involvement in host invasion, potentially serving as key players in the heightened virulence of the H race.

Recently, scientists have conducted scientific research on the study of sunflower broomrape. A. Konarska et al. (2020) examined the taxonomic attributes of the microstructure in the parasitic flowers of *Orobanche picridis*, placing particular emphasis on secretory structures. A. Krupp et al. (2019) investigated the development of phloem communication between *Orobanche cumana* parasitic plants and their sunflower hosts. A. Le Ru et al. (2021) pioneered image analysis for the automatic phenotyping of *Orobanche cumana* infection on sunflower roots. M. Abdalla et al. (2020) identified genetic variations in *Orobanche crenata* using interprost sequence repetition (ISSR) markers. R. Aly et al. (2021) examined the use of biotechnological approaches to establish crop resistance against root parasitic weeds.

Louarn et al. (2016) in their paper considered sunflower resistance to broomrape (*O. cumana*), which is regulated by specific QTL genes for different stages of parasitism. The study points to the existence of various mechanisms of quantitative resistance that control the pathogen *O. cumana* and they could be used in sunflower breeding.

The research work of Pouvreau et al. (2021) studied strigolactone-like biological activity using biological analysis of parasitic plant germination. It describes a simple and cost-effective method for evaluating the activity of compounds that exhibit strigolactone-like activity by analyzing the germination of parasitic plant seeds after they are treated with these compounds.

Xi J et al. (2022) carried out an investigation into the effectiveness of maize crop rotation coupled with *Streptomyces rochei D74* in eradicating the seed bank of *O. cumana* in agricultural areas. This research employed a dual approach involving two biological control methods to address the issue of parasitism by *O. cumana.* The findings from biological analyses affirmed that highly concentrated fermentation filtrates from *Streptomyces rochei D74* efficiently impede the germination and development of the germ tube of *O. cumana s*eeds.

In research conducted by Yang et al. (2020), a transcriptional profiling of the subterranean interaction between two divergent sunflower varieties and a root parasitic weed, *O. cumana,* was carried out. They observed that the *O. cumana* pathogen triggers insufficient protective responses in the susceptible variety compared to the resistant one, likely due to the incapacity to fully recognize the parasite’s effectors. Their study facilitated the identification of 180 proteins linked to the penetration of *O. cumana*. Functional annotation of these proteins revealed their involvement in cell wall degradation, nutrient production, and pathogenesis. An effector candidate containing the PAR1 domain within *O. cumana* had previously been identified as a potential suppressor of plant defense responses. A successful *O. cumana* infection subdued the host’s defense mechanisms, transforming the host’s roots into a source of efficient nutrient flow for the parasite.

In recent years in Ukraine, there have been no studies conducted on the racial composition of broomrape and the evaluation of resistance to it in various sunflower hybrids. The importance of these measures is determined by the need to develop a sunflower breeding strategy for broomrape and study the cellular and molecular mechanisms of crop resistance to this pathogen. To achieve effective disease control, a comprehensive understanding of the distinct mechanisms governing host protection and the pathogenesis of O. cumana is essential during the underground interaction in sunflower hybrids with varying resistance to this plant-parasitic organism. Identifying the phenological stage at which infection or incompatibility takes place is pivotal for comprehending the mechanisms of resistance to this pathogen. In the South-East of Ukraine, 5-6 races of broomrape (A-F) were identified and studied in 1990-2018 (Khablak et al., 2018). However, every year the pathogen is actively moving to the central regions of the country (Poltava, Cherkasy, Vinnytsia, Khmelnytsky, Zhytomyr regions) for those hybrids that were previously resistant and not affected. Accordingly, currently a significant territory of Ukraine is not explored. Therefore, the aim of the research was to establish the resistance of various sunflower hybrids to broomrape.

## Materials and methods

According to the research program, a vegetation experiment was conducted to determine the racial composition of the parasite and its resistance of various sunflower hybrids. The object for research in the vegetation experiment was broomrape seeds. Samples of the parasite’s seeds were collected during 2016-2020 in some of the most infected sunflower fields in the forest-steppe and Polissya.

To identify the races of broomrape, 8 sunflower hybrids from Lidea were used: ES Nirvana, ES Romantic, ES Genesis, ES Bella, ES Andrometa, ES Janis, ES Niagara, ES Artik. 20 plants of each hybrid were taken. These hybrids are resistant to A-G broomrape races. Syngenta, Arizona, Transol, Bosfora, Estrada, Kupava, Kadiks, Laskala sunflower hybrids were also used to identify races of broomrape. To determine the races of broomrape, hybrids of the Pioner sunflower selection were used: P63LL06, P64LC108 (XF 6003), P64LL125 (XF 13406), P63LE113 (XF 9026), P64HH106 (XF 13707), PR 64F66, P64LE25 (SX 9004), P64LE99 (XF 9002).

The degree of broomrape damage to plants was determined by the following scale: 7 or more brood buds per 1 affected plant (average value) (7-10 points) – strong; 4-6 brood buds (4-6 points) – medium; 1-3 brood buds (1-3 points) – weak.

Assessment of the resistance of sunflower hybrids to broomrape was carried out in soil culture using a modified method and a roll method of seed germination at the Institute of food biotechnology and genomics of the National Academy of Sciences of Ukraine (Kukin, 1960). For broomrape infection, sunflower plants were grown in soil culture in vessels with a capacity of 10 kg filled with a mixture of soil and sand in a ratio of 3:1. Broomrape seeds infected the soil mixture at the rate of 100 mg of parasite seeds per 1 kg of soil mixture. At the same time, broomrape seeds were distributed evenly in the upper third of the container. Seeds of sunflower hybrids were sown in 10 pcs. in every vessel. The plants were cultivated at 18-25°C. Maintain indoor illumination at the level of 16 hours a day in the range of 4000-7000 Lux. Watering should be carried out when the top layer of soil dries up.

30 days after sowing the seeds, the degree of damage to sunflower plants by broomrape was determined. For this purpose, sunflower plants were dug out of the containers, the root system was washed with water and the number of brood buds and broomrape seedlings on the roots was counted. Treatment of plants with preparations was not used (Fig. 1).

**Figure 1.**
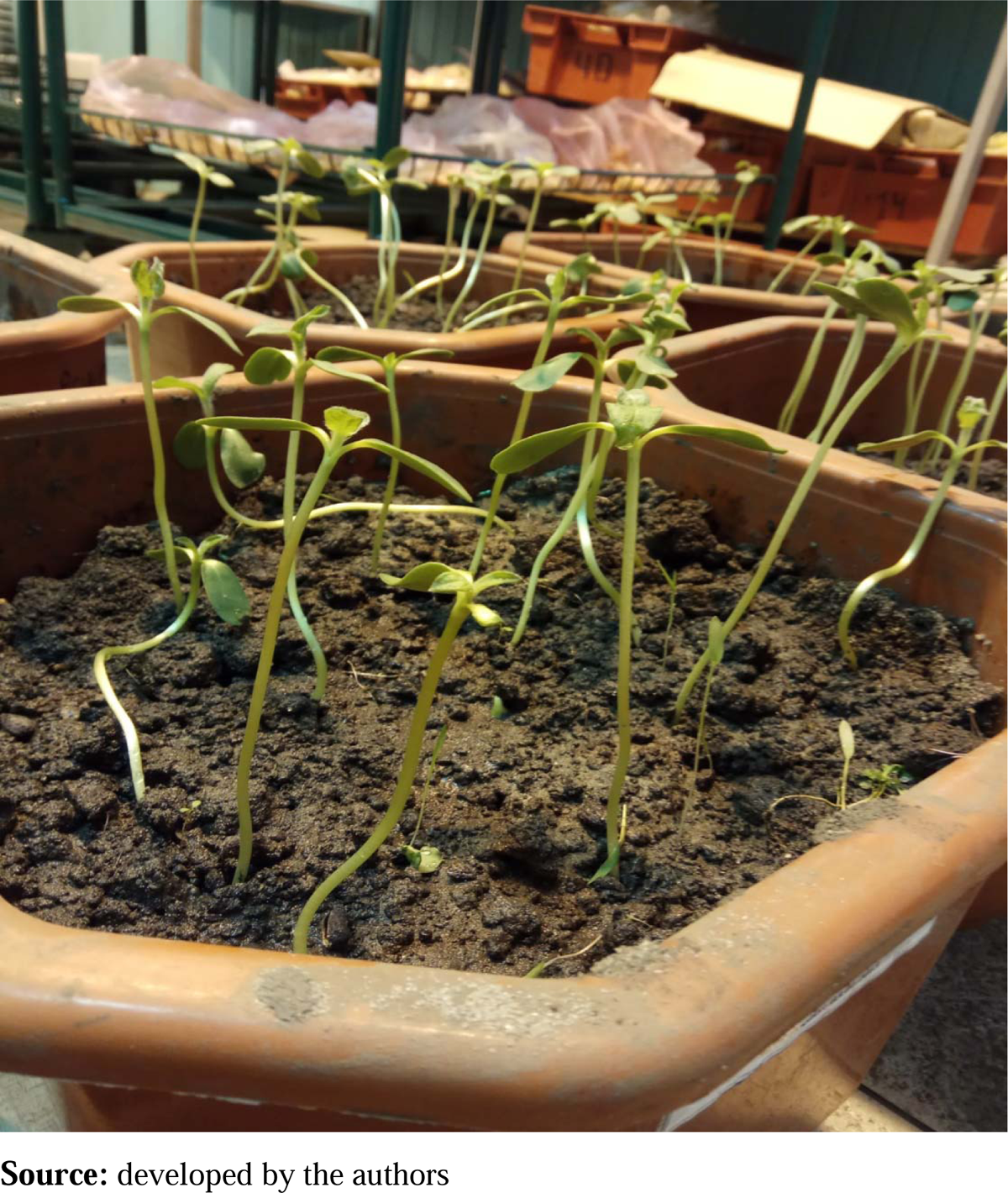
Cultivation of various sunflower hybrids in soil culture on the background infected with broomrape.

The roll method of germination of broomrape seeds consisted in the possibility of joint germination of sunflower seedlings with broomrape seeds in rolls of filter paper. The rolls were made as follows: a sheet of filter paper measuring 20 x 30 cm was folded in half to make a double sheet of 20 x 15 cm and moistened with tap water. Two-day seedlings of the sunflower hybrid were laid out so that the cotyledons went beyond the edge of the leaf, and the distance between the seedlings was 3.0-4.5 cm. There were 15 sprouts in each roll. The roots and filter paper were evenly covered with broomrape seeds. The sprouts were covered with the bent half of a sheet of paper and a roll was made. The rolls were placed vertically in a glass vessel with a small amount of water at the bottom. The vessel with the rolls was placed in an artificial climate chamber. Further co-cultivation was carried out in the artificial climate chamber “Biotron-5” produced in Ukraine for 10 days with a 16-hour photoperiod and a temperature regime of 30°C. Recording of the number of sprouted seeds was carried out on the fifth and tenth days using a stereoscopic microscope “MBS-10” produced in Ukraine.

Experimental studies of plants (both cultivated and wild), including the collection of plant material, were in accordance with institutional, national or international guidelines. The authors adhered to the standards of the Convention for the protection of biological diversity (1992) and the Convention on trade in endangered species of wild fauna and flora (1979) (Convention, 1992; Convention, 1979).

## RESULTS

Differentiation of sunflower hybrids of Lidea selection by resistance to *Orobanche cumana* Wallr. presented in Table 1.

**Table 1.**
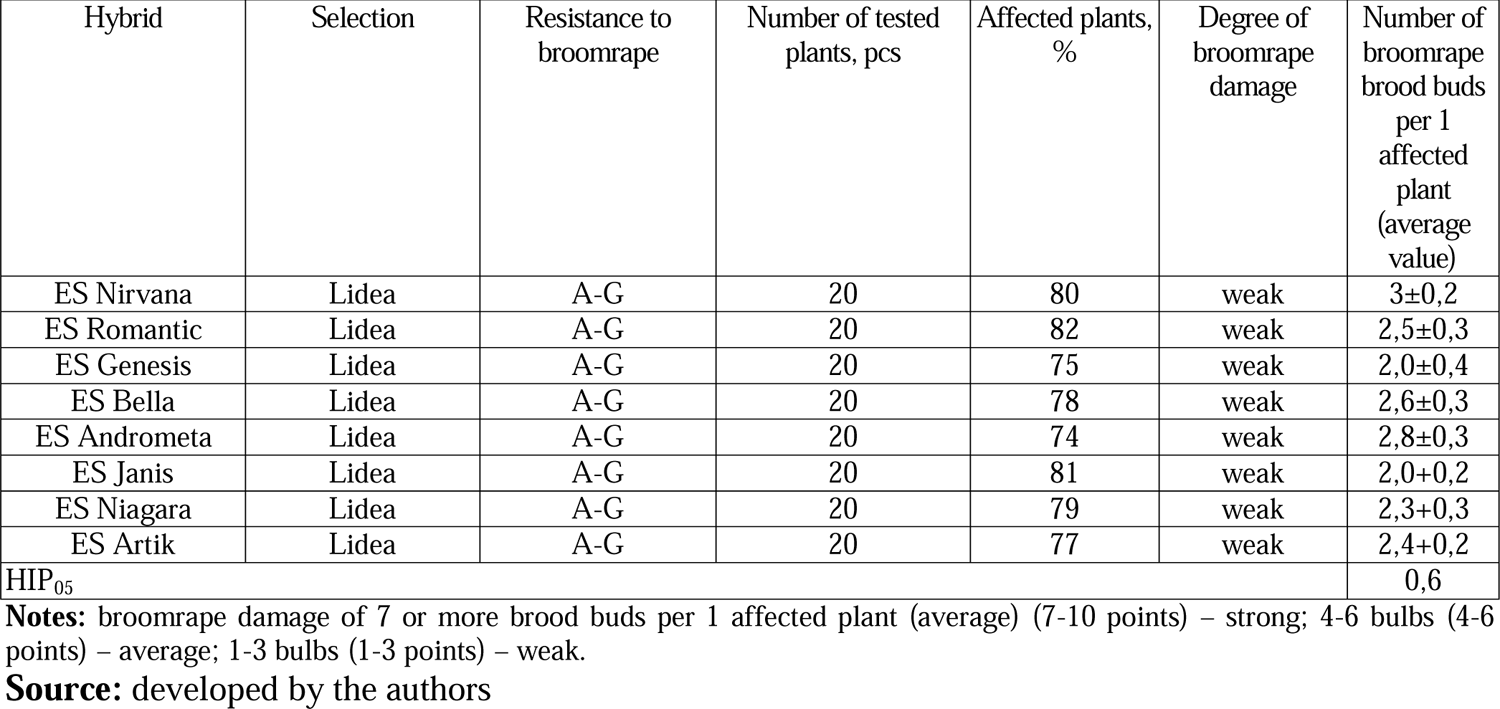
Degree of damage to sunflower hybrids by broomrape.

| Hybrid | Selection | Resistance to broomrape | Number of tested plants, pcs | Affected plants, % | Degree of broomrape damage | Number of broomrape brood buds per 1 affected plant (average value) |
| --- | --- | --- | --- | --- | --- | --- |
| ES Nirvana | Lidea | A-G | 20 | 80 | weak | 3±0,2 |
| ES Romantic | Lidea | A-G | 20 | 82 | weak | 2,5±0,3 |
| ES Genesis | Lidea | A-G | 20 | 75 | weak | 2,0±0,4 |
| ES Bella | Lidea | A-G | 20 | 78 | weak | 2,6±0,3 |
| ES Andrometa | Lidea | A-G | 20 | 74 | weak | 2,8±0,3 |
| ES Janis | Lidea | A-G | 20 | 81 | weak | 2,0±0,2 |
| ES Niagara | Lidea | A-G | 20 | 79 | weak | 2,3±0,3 |
| ES Artik | Lidea | A-G | 20 | 77 | weak | 2,4±0,2 |
| HIP<sub>05</sub> |  |  |  |  |  | 0,6 |

**Notes:** broomrape damage of 7 or more brood buds per 1 affected plant (average) (7-10 points) – strong; 4-6 bulbs (4-6 points) – average; 1-3 bulbs (1-3 points) – weak.
**Source:** developed by the authors

The results showed that sunflower hybrid plants were approximately equally affected by the parasite. Sunflower hybrids ES Nirvana, ES Romantic, ES Genesis, ES Bella, ES Andrometa, ES Janis, ES Niagara, ES Artik, tolerant to race G, were affected by broomrape. The degree of damage was not severe. On average, there were from 2 to 3 brood buds of the parasite per sunflower plant.

Sunflower hybrids with complete immunity to broomrape were not found. Since sunflower hybrids ES Nirvana, ES Romantic, ES Genesis, ES Bella, ES Andrometa, ES Janis, ES Niagara, ES Artik, resistant to race G, are affected, broomrape of races A-F (6 Race) parasitizes sunflower crops in large quantities. It is impossible to grow sunflower hybrids that are resistant to the E race of broomrape. Otherwise, it will lead to further spread of the parasite and a decrease in yield.

Due to the mild damage of sunflower hybrids resistant to race G, race H (race 8) has just begun to appear in sunflower crops. Research to identify the latter more aggressive (H and I races) parasites is complicated by the lack of sunflower resistance differentiator lines and hybrids that would allow them to be identified. Unfortunately, there are no hybrids that are resistant to race H (race 8). The best sunflower hybrids are resistant to G (race 7). In general, sunflower hybrids that are resistant to G (race 7) are tolerant to the parasite and more or less normally control broomrape. Therefore, it is recommended to grow hybrids that are resistant to 7 or higher parasite races A-G (A, B, C, D, E, F, G, H). Sunflower resistance to broomrape strongly depends on the plant’s immunity and stress state. Sunflower hybrids that are resistant to a certain race of broomrape can become more infected, having weak immunity, being in a stressful state from various stressful factors (temperature, herbicides, diseases, pests) (Fig. 2, 3).

**Figure 2.**
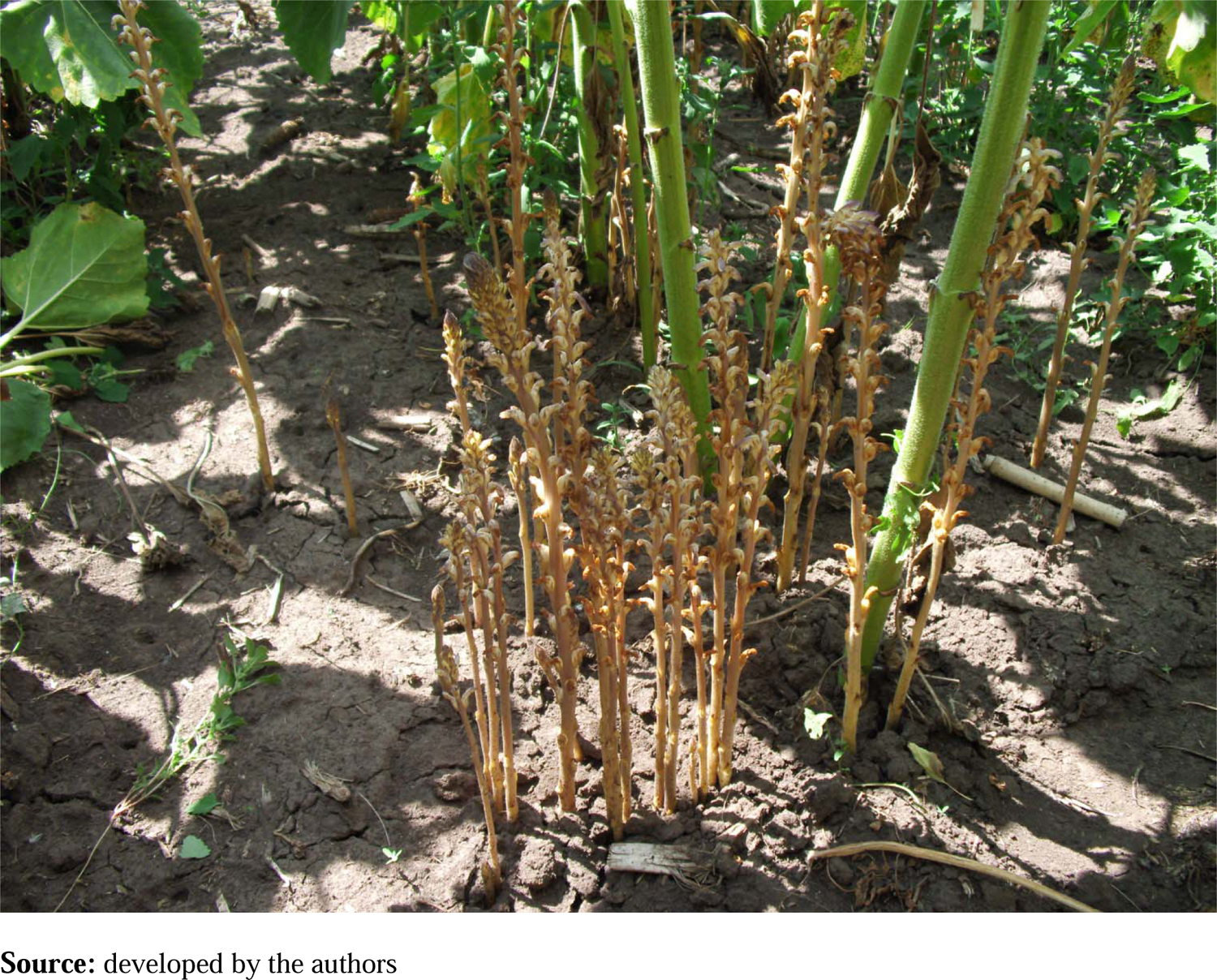
Broomrape in sunflower crops.

**Figure 3.**
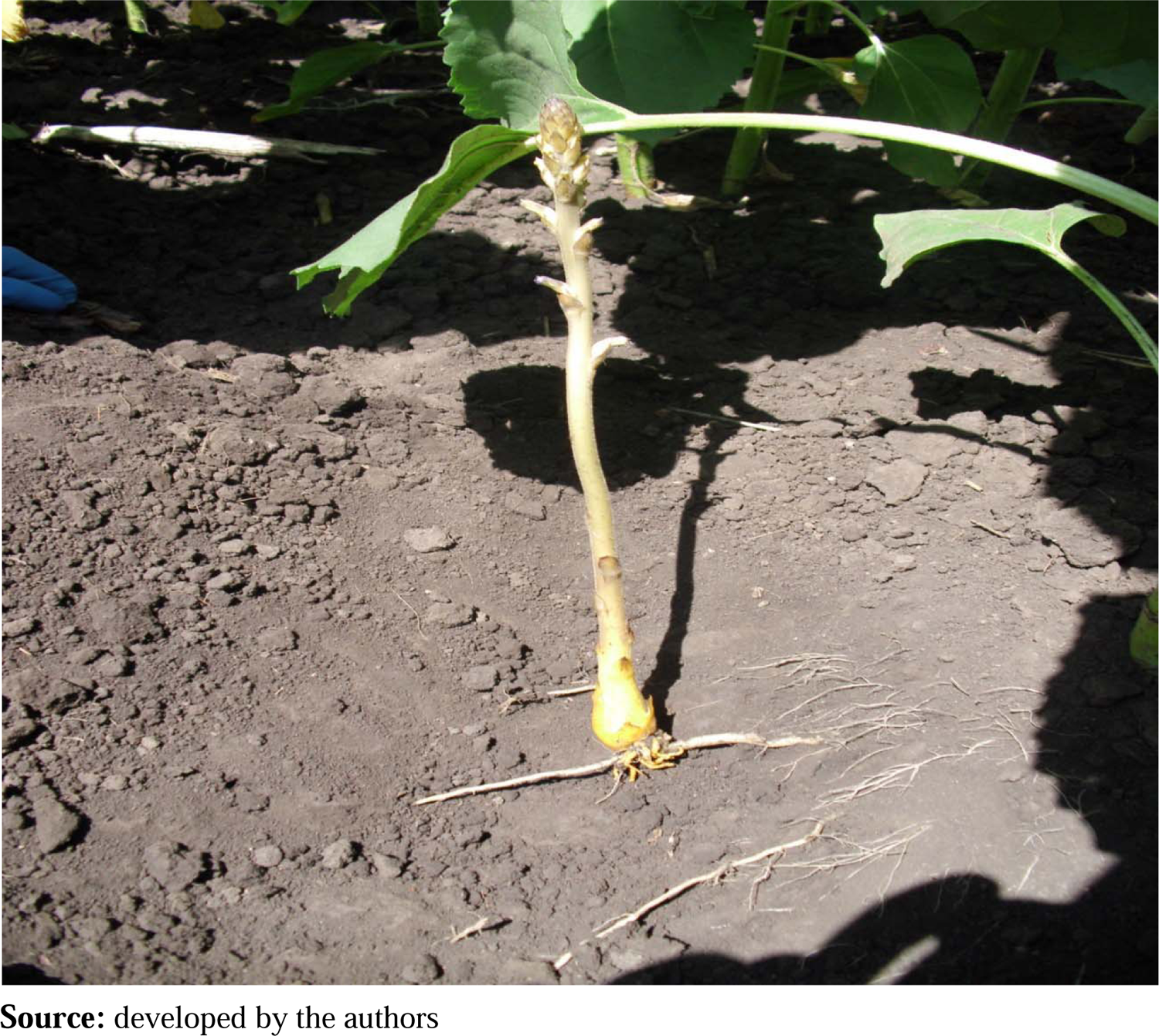
Development of broomrape on the sunflower root.

Resistance of sunflower hybrids of Syngenta selection to the parasite *Orobanche cumana* Wallr is presented in Table 2. According to the results of the research, different reactions of sunflower hybrids to the parasite were found. Sunflower hybrids Arizona, Transol, Bosfora, resistant to race F, were moderately affected by broomrape. On average, there were 5 to 6 nodules of the parasite per sunflower plant.

**Table 2.** The degree of damage to sunflower hybrids by broomrape.

| Hybrid | Selection | Resistance to broomrape | Number of tested plants, pcs | Affected plants, % | Degree of broomrape damage | Number of broomrape brood buds per 1 affected plant (average value) |
| --- | --- | --- | --- | --- | --- | --- |
| Estrada | Syngenta | A - G | 20 | 72 | weak | 2 ± 0,3 |
| Kupava | Syngenta | A - G | 20 | 78 | weak | 3 ± 0,2 |
| Arizona | Syngenta | A - F | 17 | 90 | weak | 6±0,5 |
| Kadiks | Syngenta | A - G | 20 | 74 | weak | 3 ± 0,5 |
| Transol | Syngenta | A - F | 18 | 92 | weak | 6±0,4 |
| Bosfora | Syngenta | A - F | 20 | 94 | weak | 5±0,5 |
| Laskala | Syngenta | A - G | 20 | 75 | weak | 3 ± 0,4 |
| HIP<sub>05</sub> |  |  |  |  |  | 1,3 |

**Notes:** broomrape damage of 7 or more brood buds per 1 affected plant (average) (7-10 points) – strong; 4-6 bulbs (4-6 points) – average; 1-3 bulbs (1-3 points) – weak.
**Source:** developed by the authors

Sunflower hybrids Estrada, Kupava, Kadiks, Laskala, resistant to G races, were affected to a lesser extent by broomrape. On average, there were 2-3 nodules of the parasite per sunflower plant. No sunflower hybrids with complete immunity to broomrape were found.

Since the sunflower hybrids Arizona, Transol, Bosfora, resistant to race F, were moderately affected, the sunflower crops are parasitised by broomrape of races A-G (race 7) in large quantities. Sunflower hybrids resistant to broomrape race E should not be grown. Otherwise, this will lead to further spread of the parasite and a decrease in yield.

Due to the fact that sunflower hybrids Estrada, Kupava, Kadiks, Laskala, resistant to race G, are also affected, but weakly, race H (race 8) has just begun to appear in sunflower crops. Research to identify the latter more aggressive (H and I races) parasites is complicated by the lack of sunflower resistance differentiators and hybrids that would allow them to be identified. Unfortunately, there are no hybrids resistant to race H (race 8). The best sunflower hybrids are resistant to G (race 7). In general, sunflower hybrids that are resistant to G (race 7) are tolerant to the parasite and control broomrape more or less normally. Therefore, it is recommended to grow hybrids that are resistant to 7 or more races of the parasite A-G (A, B, C, D, E, F, G, H). For example, LG 59580 is resistant to broomrape races A-G (also resistant to DuPont™ ExpressSun™ technology).

The resistance of sunflower hybrids of Pioner’s selection to *Orobanche cumana* Wallr is presented in Table 3. The results showed that the plants of sunflower hybrids were affected by the parasite in different ways. The sunflower hybrid P63LL06, tolerant to race E, was severely affected by broomrape. On average, there were 12 nodules of the parasite per sunflower plant.

**Table 3.** The degree of damage to sunflower hybrids by broomrape.

| Resistance to broomrape | Hybrid | Number of tested plants, pcs | Affected plants, % | Degree of broomrape damage | Number of broomrape brood buds per 1 affected plant (average value) |
| --- | --- | --- | --- | --- | --- |
| A-E | P63LL06 | 20 | 100 | strong | 12±0,8 |
| A-G | P64LC108 (XF 6003) | 18 | 70 | weak | 3 ± 0,3 |
| A-E + system 2 | P64LL125 (XF 13406) | 20 | 91 | average | 5±0,4 |
| A-E + system 2 | P63LE113 (XF 9026) | 15 | 92 | average | 6±0,5 |
| A-G | P64HH106 (XF 13707) | 20 | 78 | weak | 2,0± 0,4 |
| A-G | PR 64F66 | 17 | 82 | weak | 2,7± 0,3 |
| A-E + system 2 | P64LE25 (SX 9004) | 20 | 94 | average | 4±0,6 |
| A-E + system 2 | P64LE99 (XF 9002) | 20 | 90 | average | 6±0,4 |
| HIP<sub>05</sub> |  |  |  |  | 1,1 |

**Notes:** broomrape damage of 7 or more brood buds per 1 affected plant (average) (7-10 points) – strong; 4-6 bulbs (4-6 points) – average; 1-3 bulbs (1-3 points) – weak.
**Source:** developed by the authors

Sunflower hybrids P64LC108 (XF 6003), P64HH106 (XF 13707), PR 64F66, resistant to race G, were affected to a lesser extent by broomrape. On average, there were 2 - 3 nodules of the parasite per sunflower plant. The hybrids P64LL125 (XF 13406), P63LE113 (XF 9026), P64LE25 (SX 9004), resistant to race E+system 2, were infected with broomrape to an average extent. On average, there were 4-6 nodules of the parasite per sunflower plant. No sunflower hybrids with complete immunity to broomrape were found.

Since the sunflower hybrid P63LL06, resistant to race E, was severely affected, the sunflower crops are parasitised by broomrape races A-F (race 6) in large quantities. Sunflower hybrids resistant to broomrape race E should not be grown. Otherwise, this will lead to further spread of the parasite and a decrease in yield.

Due to the fact that sunflower hybrids P64LC108 (XF 6003), P64HH106 (XF 13707), PR 64F66, resistant to race G, are also affected, but weakly, race H (race 8) has just begun to appear in sunflower crops. Research to identify the latter more aggressive (H and I races) parasites is complicated by the lack of sunflower resistance differentiators and hybrids that would allow them to be identified. Unfortunately, there are no hybrids resistant to race H (race 8). The best sunflower hybrids are resistant to G (race 7). In general, sunflower hybrids that are resistant to G (race 7) are tolerant to the parasite and have sufficient control of broomrape. Therefore, it is recommended to grow hybrids that are resistant to 7 and higher races of the parasite A-G (A, B, C, D, E, F, G, H). For example, LG 59580 is resistant to broomrape races A-G (also resistant to DuPont™ ExpressSun™ technology).

The data reveals that the broomrape population parasitizing sunflower crops in the Forest-steppe and Polissya regions from 2016 to 2023 exhibits a high level of virulence, surpassing the immunity of the best foreign-bred hybrids resistant to the E, F, and G races of this parasite. The emergence of new broomrape races (E, F, G) underscores the importance of determining the parasite’s spread in sunflower crops, its racial composition, and developing measures to safeguard the crop from this pathogen across the country. To facilitate the creation of new sunflower hybrids, it is pertinent to address the challenge of developing breeding materials resistant to these novel races of the parasitic plant. Additionally, studying the cellular and molecular mechanisms of resistance to the pathogen remains relevant for further advancements in this area.

Based on the research conducted, the following strategy is proposed to protect sunflower seeds from broomrape:

1. To grow hybrids that are resistant to 7 or higher parasite races A-G (A, B, C, D, E, F, G, H). For example, LG 59580, resistant to broomrape of race A-G (also resistant to DuPont™ ExpressSun™technology).
2. In order to increase the resistance of sunflower seeds to broomrape and its immunity, to test synthetic salicylic acid preparations.
3. To grow provocative corn crops that stimulate the germination of broomrape seeds, but are not hosts of the parasite. Experiments in recent years have shown that there is a certain dependence on the corn hybrid. Research Ye et al. (2020) proved that hybrids Bilozersky 295 SV, Vityaz MV, PR38R92, DK 315) can cause germination of broomrape seeds at the level of 70%. Of course, sunflower is the best germinator, with 70% to 80% germination rate. However, it is necessary to test the corn hybrids grown in the laboratory using the roll method for the possibility of germination of broomrape seeds. After that, to recommend to grow those hybrids in the fields that cause its germination. In addition to corn, root secretions of such crops as flax, beetroot, soy, rapeseed, sorghum, Sudan grass, rapeseed and some perennial legumes (clover, sweet clover, alfalfa) can provoke the germination of broomrape seeds. But, as noted by Vurro et al. (2019), given the different root morphology, broomrape cannot attach to these crops and thus does not harm.

Gaining insights into the molecular mechanisms of host-parasite interactions holds significant promise for developing innovative and effective strategies against parasitic plants. While there has been notable progress in exploring the developmental biology of parasitic plants, the intricacies of their adaptation to various host plant species remain inadequately understood. These adaptations are evidently influenced by genetic characteristics, yet instances of rapid adaptation in genetically homogeneous populations of parasitic plants to new hosts strongly suggest the involvement of epigenetic mechanisms. A pivotal challenge lies in unraveling how host preferences are determined in diverse parasitic plants, considering the interplay between genetic and epigenetic traits. Recent strides in genomic technology offer exciting prospects for delving into the molecular intricacies of the interaction between parasitic plants and their host counterparts.

## DISCUSSION

In recent years, Ukraine has seen the spread of infection of sunflower hybrids with broomrape, which show resistance to races E, F and G. From the northern Steppe, this disease is actively spreading to the central, northern and western regions of the country due to the emergence of new races of the parasite in these areas.

This problem of harmfulness of broomrape is of global importance. Studying the parasite’s distribution area, determining its racial composition and developing measures to limit the harmfulness of broomrape are becoming urgent tasks to stabilize and increase sunflower yields, as well as to study the cellular and molecular mechanisms of crop resistance to this pathogen. Special attention should be paid to the appearance of broomrape in the central, northern and western regions of the country, its potential harm, seed germination mechanisms, identification of physiological races, integrated control and creation of sustainable sunflowers. This should contribute to the accumulation of important information for further research and effective control of broomrape.

According to current research, the best way to reduce the harmful effects of broomrape is to create sunflower hybrids that show resistance to this parasite. In order to achieve this goal, it is important to identify all races of broomrape throughout the country and build a map of the spread of the pathogen. Additionally, it is important to exchange sustainable sunflower varieties among breeding institutions in different countries. Conducting molecular studies of the interaction between Sunflower and the pathogen is useful for determining the cellular mechanisms of crop resistance to this parasite. Establishing the molecular mechanisms that regulate broomrape resistance is an important step in obtaining long-term sunflower resistance.

The examination of resistance to parasitic plants presents distinctive features when compared to dealing with other pathogens like fungi. This uniqueness arises from the close relationship between the parasite and the host, as they are relatively proximate organisms sharing numerous common morphological, physiological, and biochemical characteristics. Unlike bacteria or fungi, where many enzymes can be distinctly identified as specific to these microorganisms and differentiated from plant enzymes, the study of parasitic plants involves navigating a more intricately intertwined system of shared traits between the parasite and its host. In addition, differences in the structure of the bacterial cell wall and fungal mycelium make it easier to distinguish them in plant tissues. However, it is more difficult to consider cases of interaction between two plants that combine their tissues.

As noted by Ye et al. (2020) sunflower broomrape (*O. simapa*) shows a limited range of hosts and mainly attacks sunflowers. This species spreads across areas from Central Asia to southeastern Europe and much of northern China, leading to severe restrictions and significant losses in sunflower yields. The areas affected by this parasite are estimated at about 20,000 hectares (20-50% loss) in China, especially in the city of Bayan Nur, Inner Mongolia. Approximately 40% of sunflower fields are infected with contagion, which leads to a decrease in yield from 25% to 40%.

The research work of Shi and Zhao (2020), notes that *O. cumana* is a holoparasite plant that attacks sunflower roots. Infected sunflower plants are getting smaller, have a reduced grain-to-husk ratio, and yields are plummeting. This poses a serious threat to sunflower production worldwide. Recently, due to the import of hybrid sunflower seeds and insufficient plant quarantine, the infection has spread widely in sunflower production regions around the world.

Sunflower (*Helianthus annuus* L.) is a key oilseed that is grown all over the world for the production of butter and confectionery. Compared to other crops, such as corn, sunflower occupies an outstanding place in the successful application of heterosis. The identification of PET-1 cytoplasm, coupled with an accompanying fertility restoration system, represents a significant stride toward transforming this ornamental crop into a commercially viable oilseed crop. The primary objectives in breeding efforts encompass developing varieties characterized by high seed yield, rapid maturation, resilience to diseases (such as peronosporosis, powdery mildew, rust, necrosis, and Alternariaster leaf spot), resistance to pests (including Helicoverpa and sucking pests), and herbicide resistance. Simultaneously, there is a concerted effort to enhance the content and quality of both oil and protein. Traditional breeding methods, mutation selection, and interspecific gene transfer are employed to bolster genetic improvement and broaden the trait base. Interspecific hybridization is a key tool recognized by various research groups due to the diversity of genetic resources in wild species of *Helianthus*. The introduction of several economically important traits, such as cytoplasmic male sterility, resistance to biotic and abiotic stresses, herbicide tolerance, and improved seed quality traits, has been successful through introgression of these traits (Harbar and Knap, 2021).

Over the last two decades, remarkable strides have been made in molecular marker technologies and genomics, successfully applied in breeding to pinpoint markers associated with simple inherited traits. However, the challenge persists for breeders in addressing resistance to broomrape and other pathogens, as these traits are governed by loci of quantitative characteristics. Despite the presence of valuable genes in wild *Helianthus* species, their effective integration into the genetic makeup of varieties is hindered by the contrasting characteristics of cultivated sunflower (an annual diploid) and Helianthus species (diploid perennials with varying ploidy levels such as tetraploids and hexaploids). This necessitates the utilization of genetic engineering techniques, as highlighted in the study by Meena et al. (2022).

As noted by Rauf (2019), the cultured germ plasma of modern sunflower hybrids retains only half of the genetic diversity inherent in wild relatives of agricultural crops. The global cultivation of hybrids that share a common origin and utilize the same source of cytoplasmic male sterility poses a potential threat. Consequently, there is an imperative to leverage the existing genetic diversity within cultivated and wild germplasm to establish initial breeding lines and elite seed material characterized by high combined quality. Sunflower breeding endeavors encompass the development of breeding lines conducive to generating hybrids with heightened resistance to broomrape and other diseases, abiotic stresses, herbicide resistance, and improvements in oil content and quality. These objectives can be attained through the creation of transgenic sunflowers utilizing advanced genomic tools such as CRISPR/Cas, alongside methodologies for mapping full-genome associations. This approach facilitates the exploration of genetic modifications to enhance desirable traits and overcome challenges in sunflower cultivation.

D. Sisou et al. (2021) conducted a study of the biological and transcriptomic features of broomrape resistance in sunflowers. Visual screening and histological analysis of the roots of the confectionery sunflower variety, which is resistant to broomrape, has revealed blocking the penetration of haustoria into the roots of sunflowers. This indicates the existence of a resistance mechanism aimed at pre-establishing a haustorial interaction. Comparative RNA sequencing between samples resistant to and sensitive to broomrape has allowed revealing the genes that differed significantly in expression during infection with the pathogen. These genes encompass β-1,3-endoglucanase, β-glucanase, and ethylene-sensitive transcription factor 4 (ERF4). Historically, these genes were recognized for their association with the pathogenesis of other plant species. The findings from transcriptomic analysis, coupled with the histological investigation conducted by Sisou et al. (2021), suggest that the resistance mechanism involves the recognition of broomrape and the establishment of a physical barrier to impede the pathogen from infiltrating sunflower roots.

The downregulation of the ERF4 gene after infection should be considered in the context of the role of the endogenous hormone ethylene in controlling plant defense responses. This involves the regulation of gene expression during adaptive responses to both abiotic and biotic stresses. ERF transcription factors, unique to plants, possess a binding domain capable of interacting with the GCC-box. This specific element is present in the promoters of numerous genes associated with defense mechanisms, stress responses, and pathogenesis-related (PR) genes. Given the diversity of stresses, there is a corresponding abundance of ERF, with many of them functioning as transcription activators. For instance, AtERF1, AtERF2, and AtERF5 serve as transcription activators, whereas AtERF3 and AtERF4 function as transcription repressors for GCC-box-dependent transcription in Arabidopsis leaves (Fujimoto et al., 2000). Recent investigations by Liu et al. (2020) have identified ERF as a potential gene associated with resistance to *O. cumana* in sunflowers.

A various strategies pertaining to the sunflower’s defensive response during the initial stages of the parasite’s life cycle were outlined in studies conducted by Yang et al. (2017). These include: lignification and suberization of host cell walls. Accumulation of callose, peroxidases, and H2O2 in the cortex, along with protein crosslinking in cell walls. Increased activity of phenylalanine ammonium lyase (PAL) and elevated concentrations of phenolic compounds in host roots. Degeneration of infection following the establishment of a host-pathogen link.

When infected with sunflower broomrape, the plant’s innate immune system is involved. In this context, pathogen pattern recognition receptors (PRR), such as molecular pattern recognition receptors (PAMPs) and others (TLR), trigger activation of the hypersensitivity response and accumulation of pathogenesis-related proteins (PR). These proteins include, in particular, β-glucanases, which are PR proteins and are part of the PR-2 family. This protein family is believed to have a pivotal role in plant defense responses against pathogenic infections. For instance, in the study by Sisou et al. (2021) focusing on peas (*Pisum sativum*), it was verified that β-glucanases, with their capability to degrade the β-glucan component of the cell wall, contribute to resistance against *O. crenata*.

According to Yang et al. (2017), Pathogenesis-Related (PR) proteins are extracellular proteins that accumulate in response to pathogenic infections. Various families of PR proteins exist, including chitinases (PR-3, −4, −8, and −11), β-glucanases (PR-2), peroxidases (PR-9), and proteinase inhibitors (PR-6), which are considered among the crucial ones. Chitinases and β-glucanases play a vital role in defense by breaking down the cell walls of pathogens or releasing oligosaccharide elicitors. It’s noteworthy that chitinases do not contribute to resistance against parasitic plants due to the absence of chitin in their cell walls. Conversely, β-glucanases act on the β-glucans constituting plant cell walls, and the isolated oligosaccharides can serve as key stimulants.

The studies by Sisou et al. (2021) noted that recognition by sunflower root cell receptors of pathogen-associated broomrape molecular motifs initiates a PTI (PAMP-induced immunity or MAMP) response, which leads to suppression of ERF and suppression of PR genes, including β-glucanase. Further cleavage of the parasite’s cell walls by β-glucanase releases effectors that stimulate a second level of the plant’s immune response, known as effector immunity (ETI). As a result, a physical barrier is formed, since lignin and other phenolic compounds accumulate in the penetration zone, which leads to necrosis of the parasite, which cannot establish communication with the host vascular system.

Pathogen-associated molecular patterns (PAMPs) from the pathogen are identified by toll-like receptors (TLRs) and other pattern recognition receptors (PRRs) present in both plants and animals. This recognition mechanism enables the innate immune system to discern pathogens, thereby providing protection to the host against infections, as outlined by Soares-Silva et al. (2016).

Host recognition of a parasitic plant is crucial for resistance. As highlighted by Duriez et al. (2019), several distinct surface receptor proteins have been identified in various resistant hosts. Notably, one of these receptors, CUSCUTA 1 (CuRe1), identified in *Solanum lycopersicum*, has been demonstrated to play a significant role, albeit not the exclusive one, in conferring host resistance to Cuscuta. Interestingly, even lacking the intracellular kinase domain, CuRe1 interacts effectively with at least two SOBIR1 adapter kinases, facilitating efficient signal transmission. Similarly, in sunflower, the anticipated specific receptor for *O. cumana*, HaOr7, has been identified, characterized as a leucine-rich receptor-like kinase.

Fernández-Aparicio et al. (2022) note in their research that the OrDeb2 gene provides resistance of *Orobanche cumana* after attachment and is located in the 1.38 Mb genomic interval, which includes a cluster of receptor-like kinase and receptor-like protein genes with nine highly reliable candidates.

Thus, research to determine the spread of broomrape in sunflower crops, its racial composition, development of measures to protect the crop from this flower parasite in the conditions of forest-steppe and Polissya, and elucidation of cellular mechanisms of resistance of various *Heliánthus annuus* hybrids to the *O. cumana* parasite is an urgent task. This will allow to deepen the theory of plant immunity to pathogens, to better understand the involvement of the cell wall in plant defense reactions against pathogens, to study the mechanisms of resistance associated with the cell wall, and to understand why these mechanisms do not work when encountering some pathogens and viruses, which is fundamental for plant breeding and protection.

## CONCLUSIONS

1. From the northern Steppe of Ukraine, broomrape damage is actively moving to the central, northern and western regions of the country. The population of broomrape at the beginning of the XXI century, which parasitizes sunflower crops in the forest-steppe and Polissya, has a high degree of virulence, which overcomes the immunity of the best hybrids of foreign selection that are resistant to the G races of this parasite.
2. Sunflower hybrids ES Nirvana, ES Romantic, ES Genesis, ES Bella, ES Andrometa, ES Janis, ES Niagara, ES Artik, tolerant to race G, were affected by broomrape. The degree of damage was not severe. On average, there were from 2 to 3 brood buds of the parasite per sunflower plant. Sunflower hybrids with complete immunity to broomrape were not found.
3. Since sunflower hybrids ES Nirvana, ES Romantic, ES Genesis, ES Bella, ES Andrometa, ES Janis, ES Niagara, ES Artik, resistant to race G, are affected, broomrape of races A-F (6 Race) parasitizes sunflower crops in large quantities. It is impossible to grow sunflower hybrids that are resistant to the E race of broomrape. Otherwise, it will lead to further spread of the parasite and a decrease in yield.
4. Due to the minor damage to sunflower hybrids resistant to race G, race H (race 8) has just begun to appear in sunflower crops. Studies to identify the latter more aggressive (H and I) races of the parasite are complicated by the lack of lines-differentiators of sunflower resistance and hybrids that would allow them to be identified. Unfortunately, there are no hybrids that are resistant to race H (race 8). The best sunflower hybrids are resistant to G (race 7).
5. The emergence of highly aggressive new races of broomrape (E, F, G, and H) in the Forest-steppe and Polissya regions underscores a crucial necessity to address the challenge of developing breeding material resistant to these novel races of the parasitic plant. Resolving this issue entails a comprehensive exploration of the cellular and molecular mechanisms underlying sunflower resistance to the pathogen.
6. The results of the research revealed a different reaction of Syngenta sunflower hybrids to the parasite. Sunflower hybrids Arizona, Transol, Bosphora, resistant to race F, were moderately affected by broomrape. On average, there were 5 to 6 nodules of the parasite per sunflower plant. Sunflower hybrids Estrada, Kupava, Kadiks, Laskala, resistant to G races, were affected to a lesser extent by broomrape. On average, one sunflower plant had 2 to 3 tubers of the parasite. No sunflower hybrids with complete immunity to broomrape were found.
7. In the vegetation experiment, the sunflower hybrid P63LL06 of the Pioner selection, tolerant to race E, was severely affected by broomrape, the seeds of which were collected from fields in the central, northern and western regions of the country. On average, there were 12 nodules per sunflower plant. Sunflower hybrids P64LC108 (XF 6003), P64HH106 (XF 13707), PR 64F66, resistant to race G, were affected to a lesser extent. On average, there were 2 - 3 nodules of the parasite per sunflower plant. The hybrids P64LL125 (XF 13406), P63LE113 (XF 9026), P64LE25 (SX 9004), resistant to race E+system 2, were infected with broomrape to an average extent. On average, there were 4-6 nodules of the parasite per sunflower plant.

## THANKS

**Absent.**

